# The relict ecosystem of *Gymnosporia senegalensis* (Lam.) Loes. in an agricultural plastic sea: past, present and future scenarios

**DOI:** 10.1101/2020.04.16.044651

**Authors:** Antonio J. Mendoza-Fernández, Fabián Martínez-Hernández, Esteban Salmerón-Sánchez, Francisco J. Pérez-García, Blas Teruel, Encarna Merlo, Juan Mota

## Abstract

*Gymnosporia senegalensis* is a shrub belonging to the *Celastraceae* family, which is native to tropical savannahs. Its only European populations are distributed discontinuously along the south-eastern coast of the Iberian Peninsula, forming plant communities with great ecological value, unique in Europe. As it is an endangered species that makes up plant communities with great palaeoecological significance, the development of species distribution models is of major interest under different climatic scenarios, past, present and future, based on the fact that the climate could play a relevant role in the distribution of this species as well as in the conformation of the communities in which it is integrated. Palaeoecological models were generated for the Maximum Interglacial, Last Maximum Glacial and Middle Holocene periods. The results obtained showed that the widest distribution of this species, and the maximum suitability of its habitat, occurred during the Last Glacial Maximum, when the temperatures of the peninsular southeast were not as contrasting as those of the rest of the European continent and were favored by higher rainfall. Under these conditions, large territories could act as shelters during the glacial period, a hypothesis reflected in the model’s results for this period, which exhibit a further expansion of *G. senegalensis*’ ecological niche. The future projection of models in around 2070, for four Representative Concentration Pathway (RCP) according to the fifth report of the Intergovernmental Panel on Climate Change IPCC, showed that the most favorable areas for this species would be Campo de Dalias (southern portion of Almeria province) as it presents the bioclimatic characteristics of greater adjustment to *G. senegalensis*’ ecological niche model. Currently, these areas are almost totally destroyed and heavily altered by intensive agriculture under plastic, also causing a severe fragmentation of the habitat, which implies a prospective extinction scenario in the near future.

## Introduction

*Gymnosporia senegalensis* (Lam.) Loes. [=*Maytenus senegalensis* (Lam.) Exell] is a very thorny deciduous shrub belonging to the *Celastraceae* family, which grows up to three meters or higher. It is native to the African tropical savannahs [1–3]. Worldwide, it can be found in Africa in the southern area that expands from South Africa to Angola; throughout the east in Mozambique, Tanzania, Uganda, Kenya and Madagascar; in sub-Saharan Africa from Senegal to Ethiopia and Eritrea. It is also disseminated through the Maghreb, especially in Morocco and Algeria, as well as on the Canary Islands. In Asia, it stretches across the Arabian Peninsula, Saudi Arabia, Yemen and Oman; and seems to reach India punctually [4].

The northernmost populations are located in Europe (Spain), where it is distributed in Alicante and Malaga provinces [5,6], growing in scrubs at the most thermal areas of the southeast coastal region [7] between sea level and 300 (600) masl [8–11]. The Spanish populations seem to be adapted to a semi-arid Mediterranean climate modulated by a high proportion of dewfall from marine origin and no-frost events [12].

The presence of *G. senegalensis* in the Iberian flora dates back to the Lower Cretaceous, and is related to the palaeogeographic and palaeoclimatic history of the Mediterranean Basin [13,14]. In the last period of the quaternary, the Holocene, studies about paleoclimatic and palaeoecological trends even confirm the coexistence of this species in deciduous *Quercus* forests [15,16]. Thus, the present case represents an interesting and complex study of the distribution dynamics of a severely threatened plant species in the northernmost limit of its distribution, for which the implementation of a species distribution model could foster further understanding. In recent times, species distribution models (hereinafter SDMs) have been globally recognized as a useful tool in nature conservation and management, for instance, to refine the threat status of a species [17,18]. When applied to distribution data, they can predict distributions across geographic landscapes by multiple responses, improve image analysis or remote-sensing in order to lead the search for poorly known species [19–23], thus providing perceptions into the species’ habitat, range and abundance [24–26]. Furthermore, several authors as Elith & Leathwick [27], Benito et al. [28], Fois et al. [29] or de Luis et al. [30,31] used SDMs based on the extant localities as well as the respective current and future climate scenarios to predict the possible variation in the environmental niche of certain plant species, inferring ecological and evolutionary insights.

From a conservationist point of view, several Acts in Spain, enacted at regional and national level, currently protect this plant species [32–35], which is considered to be under vulnerable conservation status. Furthermore, this is a characteristic species for plant communities with high ecological significance [36–40] included in the Habitat Directive [41] as Priority Habitat (ANNEX I. cod. 5220* Arborescent Scrubs with *Ziziphus*). The landscape resulting from the arrangement of this vegetation in hemispherical clusters is outstanding and unique in the European context [42], in a way that is difficult to interpret, and which has resulted in different readings about its dynamics [43,44]. In addition to the landscape value, this habitat’s conservation may have positive effects that stretch beyond the characteristic community of plant species. For instance, authors as Rey et al. [45] highlight that a satisfactory management of *Ziziphus* bunches will probably lead to the conservation of many other species. Some studies show that up to 25 woody plants species present in this habitat can be gathered beneath the *Ziziphus* canopy [46], and more than 80 insect species use *Ziziphus* floral resources [47]. Wild flora has proven to be a reservoir of beneficial invertebrates for the different agricultural systems, including not only pollinators, which play a fundamental role in the functioning of ecosystems being responsible for pollinator service of numerous plant species [48,49], but also predators that might contribute positively to the biological control as an effective way to reduce pest populations surrounding orchards, vineyards or even greenhouses [50–52]. The loss of key (or minority) invertebrates consequently leads to the loss of those functions performed by them (functional diversity), which weaken and endanger the stability of these ecosystems if they cannot be replaced by other taxa [53].

For a better understanding of the palaeoecological significance of this plant species, to promote conservation measures and to identify key areas of interest, the aims of this study were to (i) model the ecological niche of *G. senegalensis* in the Iberian Peninsula using three projections in the past – Mid-Holocene (ca. 6 ka BP), LGM: Last Glacial Maximum (ca. 22 ka BP), and Interglacial Maximum (ca. 120 ka BP); (ii) evaluate the variations in the potential habitat of *G. senegalensis* modelled for the future (year 2070) in the four possible scenarios according to the representative concentration pathways (RCP 2.6, RCP 4.5, RCP 6.0, RCP 8.5); (iii) compare the retrospective and prospective results with the ecological niche model obtained from bioclimatic current data.

## Material and methods

### Study area

The study area is located in the south and southeast of Spain. More specifically, it corresponds to the coastal area biogeographically encompassed by the Baetic and Murcian-Almeriensian provinces. This territory is characterized by a Mediterranean climate, with thermo-Mediterranean thermotype, modulated from dry to semiarid ombrotype, according to the bioclimatic classification proposed by [54]. It coincides with hotspots for plant biodiversity in southern Spain in terms of rarity and endemicity [55–57], and includes semiarid as well as arid areas, which are considered among the most vulnerable ecosystems to global change drivers [58–60].

### Presence data

Presence data were collected through a wide bibliographical search combined with intensive fieldwork. Furthermore, digital sources were checked to gather distribution information on the species such as GBIF [4], ANTHOS [61] and FAME project (Threatened Flora of Andalusia Database) [62], adding details about herbaria records (HUAL, GDA, GDAC, JAEN, MGC & MUB) as well. The bibliographical data, particularly those taken from the Internet, were carefully filtered in order to only manage reliable information. All records were gathered using the WGS84 reference coordinate system; the information from databases was transformed by QGIS [63], so that it was unified. A total of 1821 georeferenced presence records of *G. senegalensis* were obtained (Fig 1 and S1 Table).

**Fig 1.**
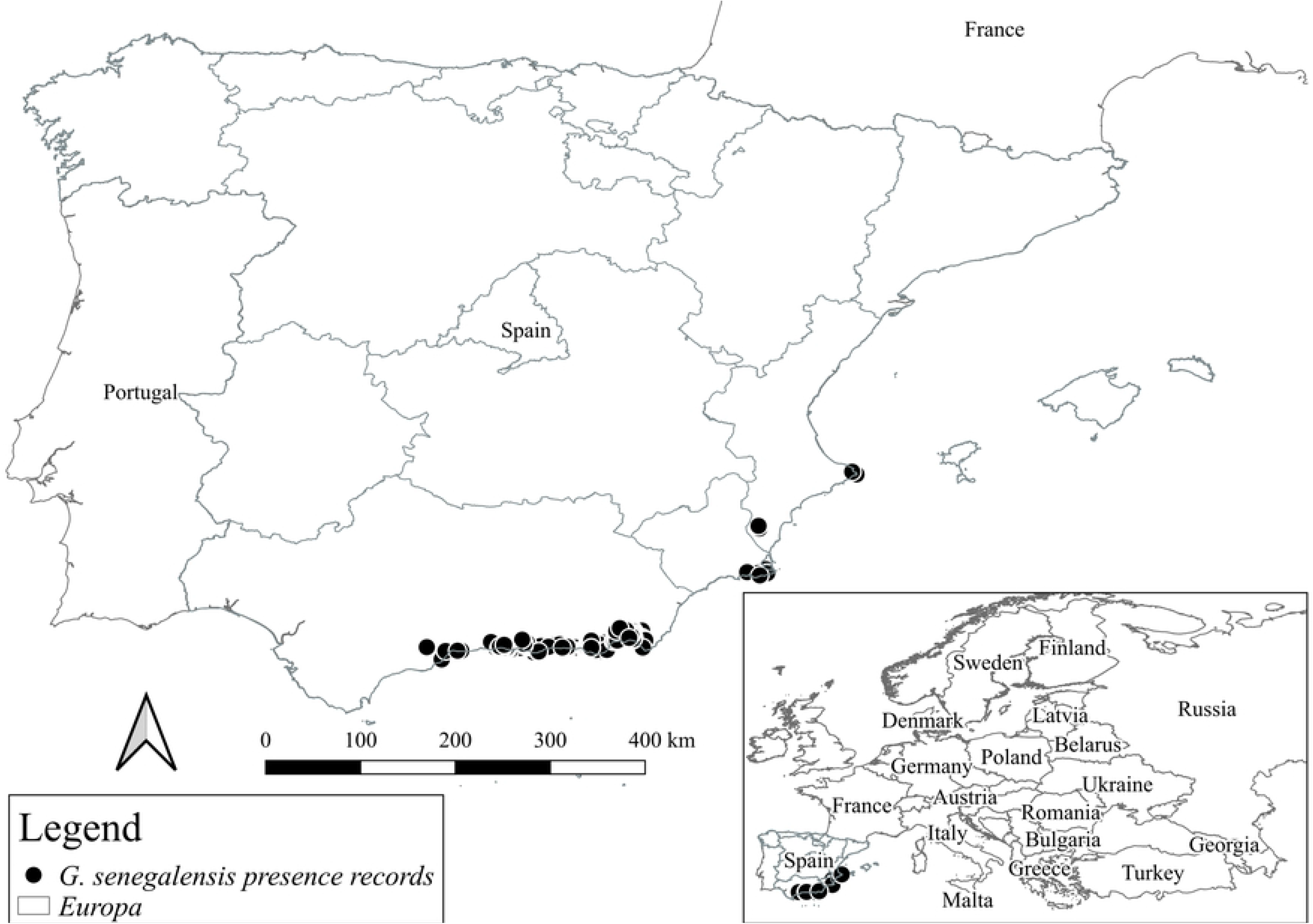
Presence records of *G. senegalensis* in Europe. The localities are represented by dots and detailed in S1 Table.

### Environmental variables

WorldClim [64] provides data on 19 bioclimatic variables considered as broadly significant from the biological point of view when modelling distribution areas [65]. Past bioclimatic data for the Mid-Holocene (ca. 6 ka BP), Last Glacial Maximum (ca. 22 ka BP), and Interglacial Maximum (ca. 120 ka BP) were used in the present study. For the current distribution modelling, bioclimatic variables were generated from the climatic data period (1960-1990). The future estimation (2070) assumed the four Representative Concentration Pathways for emissions (RCPs), according to the fifth report from the IPCC [66]. Bioclimatic data were downloaded from the Community Climate System Model (CCSM4) as the main reference for the distribution tests. This model simulates the global climate using four separate submodels for the atmosphere, earth, oceans and sea ice [67]. These bioclimatic data have a 30-second resolution, except for the Last Glacial Maximum, for which only data at 2.5 minutes were available. Each environmental variable map was adjusted to the Iberian Peninsula mask without Portugal.

To rule out the multicollinearity on the 19 bioclimatic variables, a VIF (Variance Inflation Factors) analysis was performed with the R software (S1 Fig and S2 Table) [68–70]. According to this methodology VIF value must be under 5; thus, six bioclimatic variables were selected (BIO02, BIO03, BIO08, BIO09, BIO15, BIO19) to execute the modelling process (Table 1).

**Table 1.** Modeling process variables.

|  |  |
| --- | --- |
| BIO02 | Mean Diurnal Range (Mean of monthly (max temp - min temp)) |
| BIO03 | Isothermality (BIO2/BIO7) ($\times 100$) |
| BIO08 | Mean Temperature of Wettest Quarter |
| BIO09 | Mean Temperature of Driest Quarter |
| BIO15 | Precipitation Seasonality (Coefficient of Variation) |
| BIO19 | Precipitation of Coldest Quarter |

Bioclimatic variables (Bi) selected in the VIF analysis, from the 19 variables available on WorldClim website.

### Species distribution modeling (SDMs)

MaxEnt 3.3.3 [71] was used to model the potential habitat of *G. senegalensis* in the different proposed scenarios. The application of the Maximum Entropy principle to estimate ecological niche modelling and potential distribution areas follows the studies of Phillips et al. [72] and Phillips & Dudík [73]. Said software has been used in many fields of science proving its validity [74–77], being widely recognized as the most used tool, even for small sample sizes and poorly known species’ distributions [24]. The modelling process is able to indicate suitable conditions for species in areas where their presence has not been registered or evidenced; several studies supported the reliability of SDMs [21]. Since this process is iterative, the modelling outcomes can highlight other factors related to the physical environment, anthropogenic influences or conditions of growth and reproduction of the species [78–80].

MaxEnt can operate with information about presence, which often represents the largest set of available data [81]. Only-presence-data algorithms usually represent the spatial distribution of the fundamental ecological niche of a species, while algorithms based on presence-absence data characterize more closely the distribution of the effective ecological niche [82]. The present study has been conducted with presence data alone, since a presence-absence modelling might generate controversies in light of the recent dynamics of the largest *G. senegalensis* population. Land changes due to the evolution of intensive crops in the province of Almeria [83], which may result in defining an area with optimal characteristics for this species as an area of absence, could offer an incorrect outcome and decrease biological significance in the interpretation of the model [84,85].

In the MaxEnt configuration, 1000 was established as the maximum iteration parameter; the convergence limit was set at 0.00001 with a value of 0.001 for regularization [72]. Hinge and threshold functions were disabled, leading to easily interpretable response curves and further adjustment to the Ecological Niche Theory [86]. The program produced a set of maps in which each pixel represents a value between 0 and 1, being thus interpreted as an index of habitat suitability. It also allowed for the generation of response curves for the species in relation to the variables used, noting their suitability, the evaluation of optimal values, the tolerance intervals and the various variables’ thresholds.

### Potential distribution of *G. senegalensis* in Europe

MaxEnt models’ results for *G. senegalensis* were adjusted by comparing its distribution with the presence of species that characterize different habitats together with [11,83,87], such as *Periploca angustifolia* Labil. (they coexist in the association *Mayteno europaei-Periplocetum angustifoliae* Rivas Goday & Esteve; *Ziziphus lotus* (L.) Lam, (both species subsist in the association *Gimnosporio europaei-Ziziphetum loti* F. Casas), or mesophilic species such as *Buxus balearica* Lam. and *Cneorum tricoccon* L. (the three coexist in the association *Cneoro tricocci-Buxetum balearicae* Rivas Goday & Rivas Mart.). Chorological data for these plant species were obtained from GBIF and ANTHOS databases (op. cit). In addition, the current final map that describes the potential habitat of *G. senegalensis* and its decrease in Europe were designed as contiguous areas by combining presence records, species modelling, and photo-interpretation of polygons performed in the Spanish SIOSE project [88] as was done by Mendoza-Fernández et al. [89].

## Results and discussion

### Potential habitat models in the past

The model obtained for the Last Glacial Maximum (LGM) showed a significant dominance of the *G. senegalensis* habitat towards the southeast and the east of the Iberian Peninsula (Fig 2 and S2 Fig), exhibiting high suitability in areas nowadays submerged, and along the coastal plains of Almeria province. Probably, the populations were more stable in said area, acting as a genetic diversity reservoir in the glacial period. Médail and Diadema [90] identify the existence of such shelters as a crucial event in a context of global change. Furthermore, since the LGM possibly represented the most favorable scenario for this plant, the idea that *G. senegalensis* could not be as xerophyte an element as considered may be reinforced. During this period, weather conditions became more humid than in the Mid-Holocene and the Interglacial Maximum. In addition, in the case of the southern Iberian Peninsula, the temperature decrease was not as extreme as in other parts of Europe, favoring a climate characterized by relatively mild temperatures and intense rainfall [91]. On the contrary, the resulting model for both the Mid-Holocene and the Last Interglacial periods did not show such habitat suitability for the plant. While the results for both periods were quite similar, it should be noted that the Mid-Holocene scenario presented a medium suitability for this plant since the modelling process was very strict with the bioclimatic variables developed for this time interval. The outcome of this period subsequent to the LGM, where the niche model showed the greatest habitat amplitude for *G. senegalensis* in the southeast of Spain, may be understood as the consequence of a delay in the dynamics of the *G. senegalensis* populations from the LGM long-lasting the Mid-Holocene period, as previous studies in Sierra de Gador suggest [16], which shows the gradual decline of *G. senegalensis* and other deciduous *Quercus* genus species that might have been more widely distributed during the LGM.

**Fig 2.**
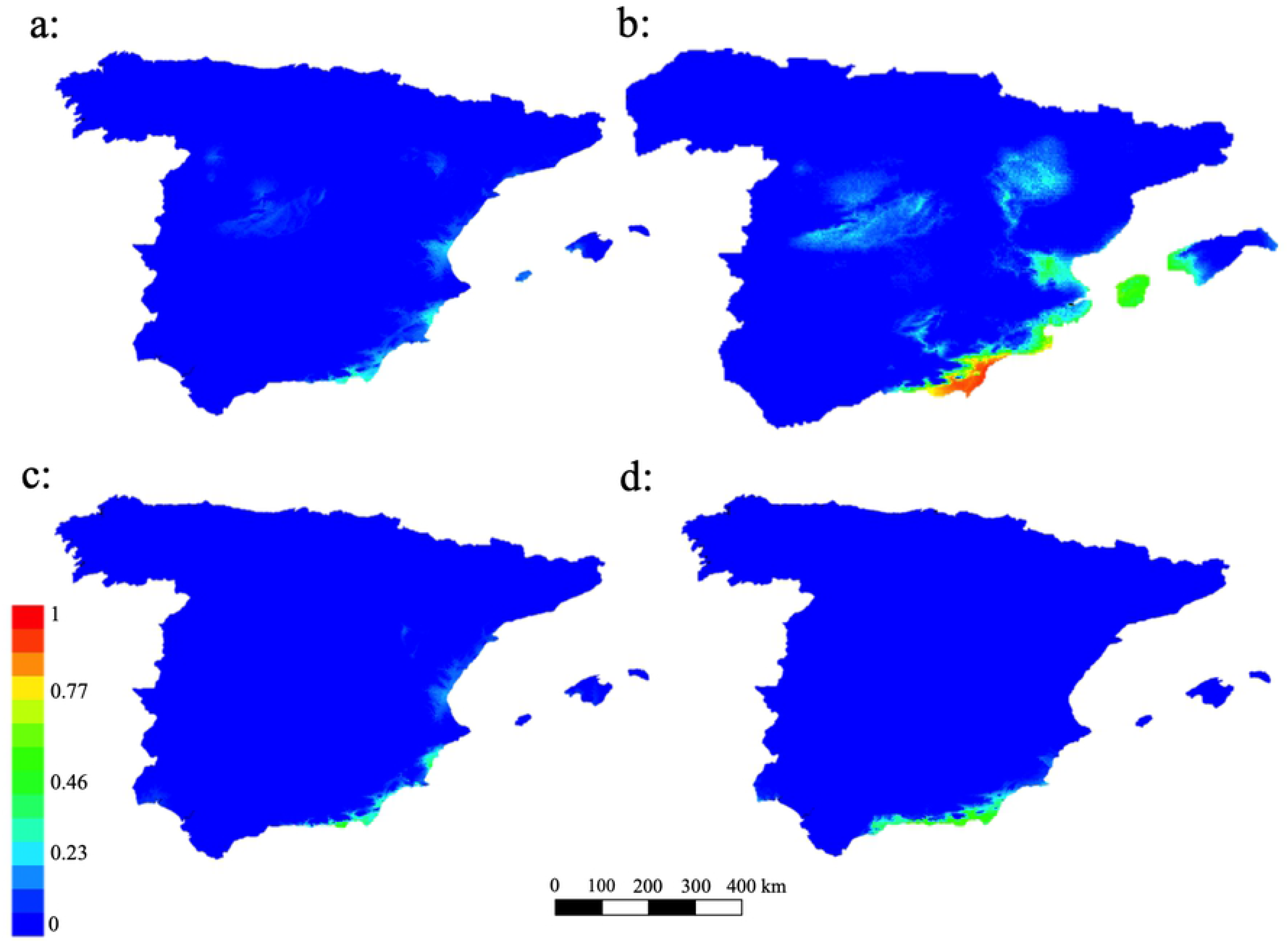
Past models for *G. senegalensis*. Plots of potential habitat of *G. senegalensis* in the past – a: Interglacial Maximum (ca. 120 ka BP); b: Last Glacial Maximum (ca. 22 ka BP) Mid-Holocene (ca. 6 ka BP)– and d: Present.

### Models for the potential habitat towards the future (2070)

The results of the future models with a projection for the year 2070 showed a slight variation in suitability among them, in which neither considerable habitats increase nor a significant decrease of it was observed. In the different scenarios, the zones exhibited small geographical variations, which slightly expand or reduce *G. senegalensis*’ ecological niche (Fig 3). However, habitat suitability was not altered, remaining permanent, with intermediate values in the four models. The results reinforce the thesis of *G. senegalensis*’ mesophilic behavior, and contrary to expectation, the global temperature increases projected in the different RCPs scenarios could predict the probable deterioration of the habitat. Nevertheless, a specific area was highlighted, the coastal plain in the south of the province of Almeria (from sea level to 350 masl approx.), where the four models corresponding to each RCPs scenario remained constant in terms of suitability, and offered the maximum values for all models. This fact makes this area a very probable one where potential *G. senegalensis* habitat is predicted in any of the 2070 scenarios. In addition, it supports the idea that this would be the most suitable area for the plant, in light of the four possible futures with an irradiative effect increase. Despite the fact that this area might be treated in the short term as one of the few habitats in the European continent available for this plant, it is currently a very altered territory, especially due to land-use changes resulting from the proliferation of intensive agriculture, which has reached up to 90% of the potential area of this species [83,89,92,93].

**Fig 3.**
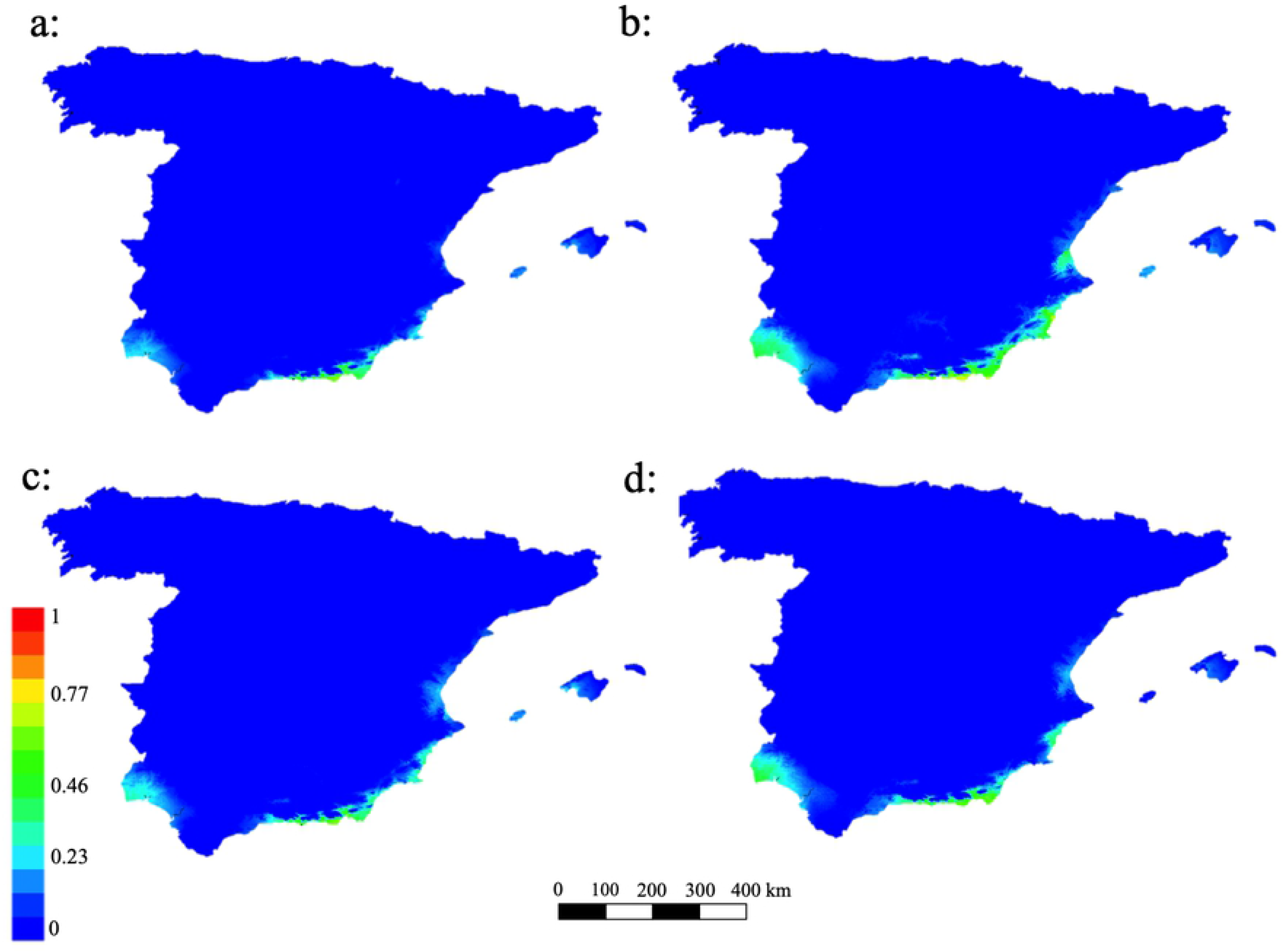
Future models for *G. senegalensis*. Plots of the potential habitat projected into the future according to the representative concentration pathways a: RCP 26; b: RCP 45; c: RCP 60; d: RCP 85.

### Current extent of occurrence in Europe

The results achieved by combining presence records, species modelling, and photo-interpretation of polygons demonstrated that *G. senegalensis*’ current extent of occurrence (EOO) in Europe is approximately 166721.2 ha, less than 47% of the suitable potential area (Table 2 and Fig 4).

**Table 2.** EOO of *G. senegalensis* in Spanish political divisions and Europe.

|  | Available EOO (ha) | Suitable area (ha) | Remaining EOO (%) |
| --- | --- | --- | --- |
| **Andalucia** | 149952.1603 | 284747.7731 | 52.6614 |
| **Murcia** | 11079.9679 | 32266.6956 | 34.3387 |
| **Valencia** | 5689.0772 | 31891.6983 | 17.8387 |
| **Europe** | 166721.2055 | 348906.1669 | 47.7840 |

Values extracted from the detailed map of land uses of Spain (SIOSE).

**Fig 4.**
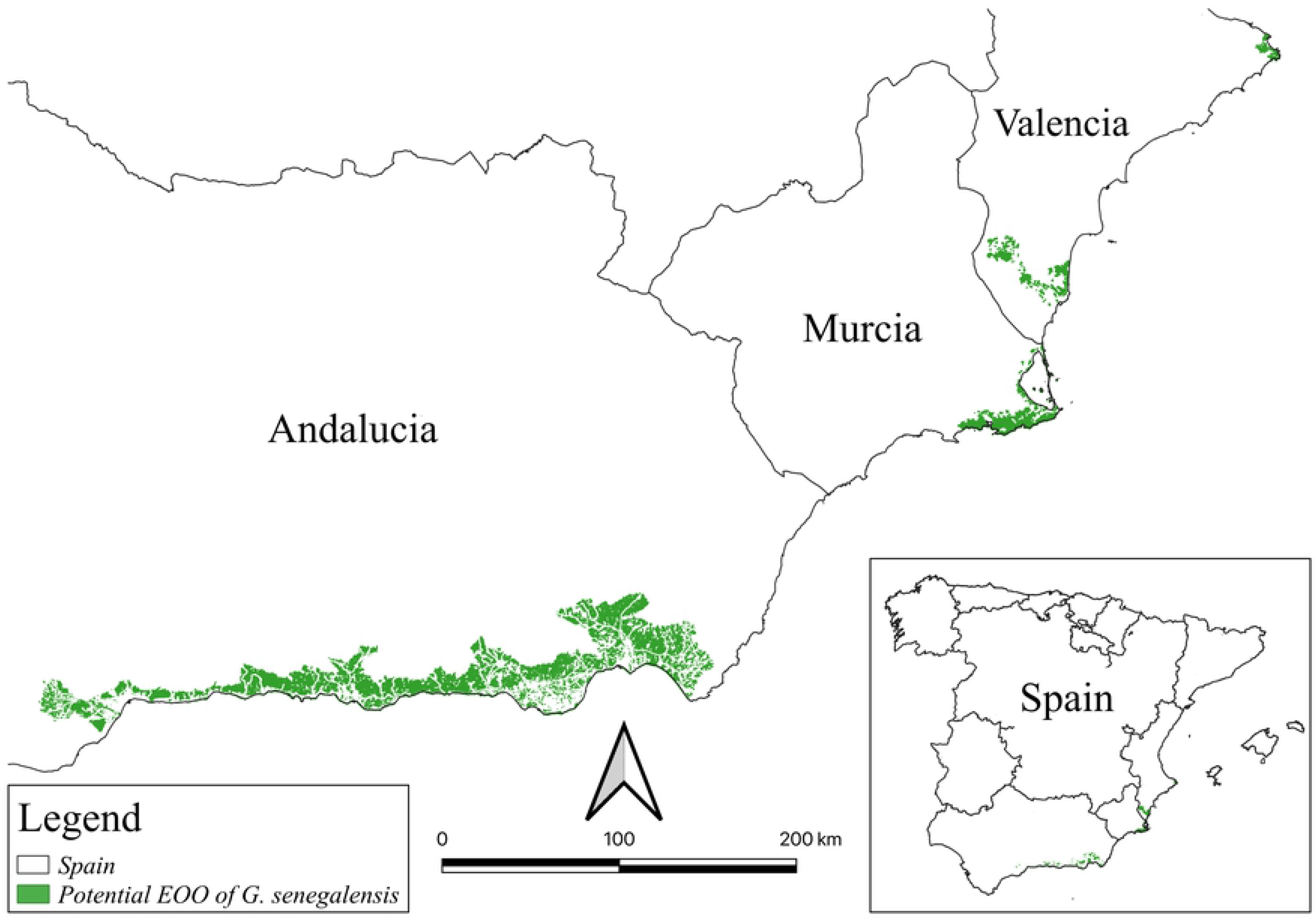
Potential EOO *G. senegalensis*. In green color polygons of the available habitat.

As can be observed in Fig 1, there are no presences of *G. senegalensis* in the east coast of Andalucia (province of Almeria) as well as in a significant part of the Murcian coast. This is an important point to take into consideration when interpreting the modelling results, since this area is the driest in the Spanish southeast, but where *Z. lotus* and *P. angustifolia* plant communities develop and grow. Unifying the distribution of the aforementioned species in a single map helped to realize that *G. senegalensis* exhibits a less xerophytic behavior than *Z. lotus* and *P. angustifolia*, since the former does not coexist with the latter in the Murcian-Almeriensian bordering zone, thus proving *G. senegalensis*’ more mesophilic character (S3 Fig).

The slight probability of presence recorded to the north and west of Spain may be caused by common temperature variables or the coincidence of the summer drought period, related to the savannoid character of this plant, since there is no evidence of any past or current presence in these areas.

In order to achieve higher accuracy in establishing the influence of bioclimatic scenarios, models of singular thermophilic and xerophilous species that coexist with *senegalensis* could be compared in equal space-time conditions [91]. Hence, several models from different communities may be obtained and thus their spatial-temporal dynamics analyzed. In addition, indirect gradient variables corresponding to the physical characteristics of the territory (elevation, orientation, slope, geology, etc.) could be included in the model (at least at present and future predictions), since they might show a good correlation with the patterns of species distribution by combining resource gradients and direct gradients [94].

## Conclusions

1. From a palaeoecological point of view, the glaciation conditions favored the extension of the potential niche for *G. senegalensis* in the Iberian southeast, demonstrating that this plant does not have as xerophytic a behavior as *Z. lotus* or *P. angustifolia*, since the LGM model is the one that presents the highest suitability. This fact may be related to an increase in humidity in that period [95]. This conclusion is consistent with two other types of evidence:

a. the presence of *G. senegalensis* growing and showing its maximum abundance in deciduous forests located in Sierra de Gador during the Mid-Holocene according to palynological studies,
b. the link between *G. senegalensis* with *C. tricoccon* and *B. balearica*, which have mesothermophilic characteristics, capable of generating debate on the *Gymnosporia*-*Periploca*-*Ziziphus* trilogy, which was considered until now as a set of Ibero-African species with a hyperxerophytic conduct. This could explain the absence of *G. senegalensis* from Cabo de Gata to the Murcia region, being this area the most arid and with the smallest rainfall in the entire southeast, where conversely both *Z. lotus* and *P. angustifolia* are frequent.
2. In the near future, still affected by Global Change, the southern part of the province of Almeria may become the most suitable area for *G. senegalensis* in the Iberian Peninsula. However, the natural area available for this plant’s development might be limited by the proliferation of intensive crops, which already occupy more than 90% of the habitat in this zone. This would cause a serious conservation problem for this species and increase its extinction risk due to massive occupation, fragmentation and land-use changes.
3. Although the area known as Campo de Dalias (southern portion of Almeria province) has suffered the greatest habitat loss in terms of the extent of occurrence and occupancy area, this is a key territory for the future of the species, as demonstrated by the distribution models generated throughout this research. Therefore, safeguarding the last remaining redoubts of the Spanish *G. senegalensis* communities here is considered absolutely essential.
4. In addition to conserving some semi-natural stronghold for this species formation in the aforementioned area, there are two additional strategies that may favor its preservation. The first one relates to the use of this species with the aim to establish hedges, vegetal fences [44] or small clusters that serve as a refuge for pollinators and predatory species of insects or other harmful pests for crops. In general, insects are considered one of the most effective animal groups in the pollination processes, thus their presence in natural systems is essential. Nevertheless, intensive agriculture and continuous landscape modification through land-use changes are associated with the decrease in the population of pollinators [96], and/or severe alterations in their population concentrations [97]. Moreover, the agricultural use of pesticides and broad-spectrum chemicals produces a decrease not only in pest insects but also in all the entomofauna associated with these systems, thus affecting the insects of less altered neighboring habitats [98]. The combination of *Gimnosporia* and *Ziziphus* bunches could considerably improve this strategy, and hence favor a more sustainable development model in an area where highly intensive agriculture production can be understood as a industrial system if inputs and residues are attended to [83].

The second strategy of great interest would be the creation of green spaces, by means of peri-urban parks, where some of the remaining fragments of this community that cannot be conserved within a legally protected area, may be integrated.

## Acknowledgements

This study has been made possible through the projects ‘CEIJ-012 Integrated study of coastal sands vegetation (AREVEG)’ and ‘CEIJ-009 Integrated study of coastal sands vegetation (AREVEG II)’ sponsored by CEI·MAR; ‘Assessment, Monitoring and Applied Scientific Research for Ecological Restoration of Gypsum Mining Concessions (Majadas Viejas and Marylen) and Spreading of Results (ECORESGYP)’ sponsored by the company EXPLOTACIONES RÍO DE AGUAS S.L. (TORRALBA GROUP) and ‘Provision of services, monitoring and evaluation of the environmental restoration of the mining concessions Los Yesares, María Morales and El Cigarrón’ sponsored by the company Saint Gobain Placo Iberica S.A. We thank Beatrice Antolin for reviewing the English translation of the text.

## Supporting information

**S1 Fig. Results of VIF analysis.** Cluster analysis of the 19 bioclimatic variables (bi_sp) used in the VIF analysis. Abbreviations are listed in Table S2.

**S2 Fig. Results of gains and Roc curve.** MaxEnt analysis of the variables.

**S3 Fig. Distribution of** *Z. lotus*, *C. Tricoccon*, *B. Balearica* **and** *P. angustifolia*. Records at 10km^2^in south and east of Spain.

**S1 Table. Presence records of** *G. senegalensis*. Geographic coordinates of presence records gathered form several sources.

**S2 Table. Bioclimatic variables (Bi).** Variables available at WorldClim website.

